# Regulation of Ferroptosis in Obesity: Muscle Type-Specific Effects of Dietary Restriction and Exercise

**DOI:** 10.1101/2024.08.04.605473

**Authors:** Fujue Ji, Haesung Lee, Jong-Hee Kim

**Affiliations:** Department of Physical Education, College of Performing Arts and Sport, Hanyang University, 222 Wangsimni-ro, Seongdong-gu, Seoul, Republic of Korea; BK21 FOUR Human-Tech Convergence Program, Hanyang University, 222 Wangsimni-ro, Seongdong-gu, Seoul 04763, Republic of Korea

**Keywords:** Obesity, Ferroptosis, Muscle type-specific, Dietary restriction, Exercise

## Abstract

**Background:** Obesity is a significant global health issue and a risk factor for numerous diseases. Ferroptosis, an iron-dependent regulated cell death, is triggered by iron overload and the excessive accumulation of lipid peroxidation mediated by reactive oxygen species. Studies has identified a strong association between ferroptosis and obesity. Additionally, dietary restriction (DR) and DR combined with exercise (DR+Ex) are effective strategies for managing obesity and ferroptosis. However, the regulation of ferroptosis and its signaling pathways in skeletal muscle under conditions of obesity, DR, and DR+Ex remains poorly understood.

**Methods:** Mice were divided into four groups: normal diet, high-fat diet, high-fat DR, and high-fat DR+Ex. All mice were fed ad libitum with either a normal or high-fat diet for the first 14 weeks, followed by the respective interventions for the subsequent 8 weeks. Mice muscle ferroptosis were examined by immunohistochemistry, Hematoxylin & Eosin, Masson’s trichrome, Prussian blue staining, and Western-Immunoblot.

**Results:** The high-fat diet resulted in increased inflammatory cell infiltration, fibrosis, and iron accumulation. Red and white muscle showed increased expression of 4-HNE, regulated by GPX4 and NCAO4, respectively. DR and DR+Ex reduced downstream 4-HNE expression by regulating GPX4 in red muscle.

**Conclusion:** Due to metabolic differences, obesity-induced ferroptosis in skeletal muscle and the regulation by DR+Ex exhibiting fiber type-specificity. Specifically, red and white muscle respond to obesity-induced ferroptosis through different pathways; red muscle can inhibit obesity-induced ferroptosis through DR+Ex by GPX4. This deepens the understanding of mechanisms related to skeletal muscle cell death and provides scientific data support for the personalized treatment of related diseases.

## 1. Introduction

Obesity has emerged as a significant global health issue, affecting millions of individuals worldwide. Its detrimental effects on human health are well-documented, with numerous studies linking it to chronic diseases such as cardiovascular disorders, type 2 diabetes, and certain types of cancer (Blüher, 2019). Recent research has found that various chronic diseases caused by obesity are related to its induction of multiple cell death pathways, including ferroptosis (Stockwell et al., 2017).

Ferroptosis is an iron-dependent form of regulated cell death caused by iron overload and the excessive accumulation of lipid peroxidation products mediated by reactive oxygen species (ROS) (Dixon et al., 2012). This unique cell death mechanism has been implicated in the pathogenesis of various diseases across different organs and tissues, including neurodegenerative disorders, liver diseases, and kidney injury (Stockwell & Jiang, 2019). Studies have found that obesity can lead to tissue or organ dysfunction and damage by regulating ferroptosis. Specifically, obesity regulates cellular ferroptosis by modulating iron, lipid, and amino acid metabolism signaling pathways, thereby inducing and worsening tissue or organ dysfunction and damage (He et al., 2023).

Skeletal muscle is the largest metabolic organ in the human body, playing a crucial role in energy metabolism and fat oxidation, and is thus an indispensable part of overall metabolic health (Franco-Romero et al., 2024; Mizushima, 2007). Studies have found that obesity can lead to skeletal muscle dysfunction and even diseases, such as contractile dysfunction and sarcopenic obesity (Collins et al., 2018; Kalinkovich & Livshits, 2017). Furthermore, ferroptosis-related signaling has been identified in diseases such as sarcopenia, rhabdomyolysis, rhabdomyosarcoma, and fatigue due to excessive exercise (Ding et al., 2021; Guerrero-Hue et al., 2019; Huang et al., 2021). These skeletal muscle diseases severely impact patients’ quality of life and physical function, ultimately increasing mortality rates. These studies indicate that skeletal muscle health is closely related not only to obesity but also to ferroptosis. However, it remains unclear whether ferroptosis is involved in the negative regulation of skeletal muscle health by obesity. Additionally, skeletal muscle is composed of different muscle types (white and red), which exhibit distinct metabolic characteristics, and their responses to ferroptosis under obesity-induced stress are also unclear. Addressing these scientific questions will help us further understand the negative effects of obesity on skeletal muscle and provide new therapeutic strategies for treating related diseases in the future.

Dietary restriction (DR) has been extensively studied for its potential health benefits, not only ameliorates obesity-induced skeletal muscle dysfunction (Suga et al., 2014) but also regulated the signaling pathway associated with ferrpotosis (Fang et al., 2021). Exercise (Ex) is widely recognized for its positive impact on health (Ruegsegger & Booth, 2018), not only improves obesity-induced skeletal muscle dysfunction (Ostler et al., 2014) but also modulates the expression of ferroptosis signaling in the brain, kidney, and nerves (Chen et al., 2023; Liu et al., 2022; Xiang et al., 2023). Both CR and Ex demonstrate dual regulatory effects on obesity and ferroptosis. Moreover, dietary restriction combined with exercise (DR+Ex) is acknowledged as a method that more effectively counteracts the negative effects of obesity and lipid peroxidation than DR or Ex alone (Khachatoorian & Samara, 2018; Kim et al., 1996; Kvedaras et al., 2020; Li et al., 2017). However, the regulatory effects of DR and DR+Ex on obesity-induced ferroptosis in skeletal muscle remain unclear. Addressing this gap can provide scientific data to support the development physical therapy for obesity- and ferroptosis-related skeletal muscle diseases in the future.

Therefore, this study will first reveal the regulatory mechanisms of obesity on ferroptosis in skeletal muscle. Subsequently, it will investigate the regulatory mechanisms of CR and CR+Ex on skeletal muscle ferroptosis induced by obesity. Understanding the molecular mechanisms underlying these processes could provide valuable insights into the development of targeted interventions for obesity-related muscle disorders and promote overall health and well-being.

## 2. Methods

### 2.1 Animals and Experimental Design

Male C57BL/6N mice (14-month-old) were randomly divided into four experimental groups: Normal diet (ND, n=7), High-fat diet (HF, n=7), 20% high-fat dietary restriction (HFR, n=7), and 20% high-fat dietary restriction with voluntary wheel running exercise (HFRV, n=7). For the first 4 months, all mice were fed either a standard chow diet (ND; Teklad Global 2018, Envigo Inc.) or a high-fat diet (HF; 45% kcal from fat, D12451, Research Diets Inc.) (Table 1). In the subsequent 4 months, dietary restriction (DR) and dietary restriction combined with exercise (DR+Ex) interventions were applied to the respective groups (Figure 1). The mice were housed in a controlled environment, maintained at a temperature of 22 ± 2 °C with 50–60% humidity, and subjected to a 12-hour light/dark cycle. All animals had ad libitum access to water and were allowed physical activity. The experimental procedures were approved by the Institutional Animal Care and Use Committee (IACUC) of Hanyang University (HYU 2021-0239A).

**Table 1.** Diet composition.

| Group | ND |  | HF, HFR, and HFRV |  |
| --- | --- | --- | --- | --- |
|  | Teklad Global 2018, Envigo |  | D12451, Research Diets |  |
| Diet | Inc. |  | Inc. |  |
|  | gm% | kcal% | gm% | kcal% |
| Protein | 18.4 | 30 | 24 | 20 |
| Carbohydrate | 44.2 | 50 | 41 | 35 |
| Fat | 6 | 20 | 24 | 45 |
| Total |  | 100 |  | 100 |
| Kcal/gm | 3.1 |  | 4.73 |  |

**Figure 1.**
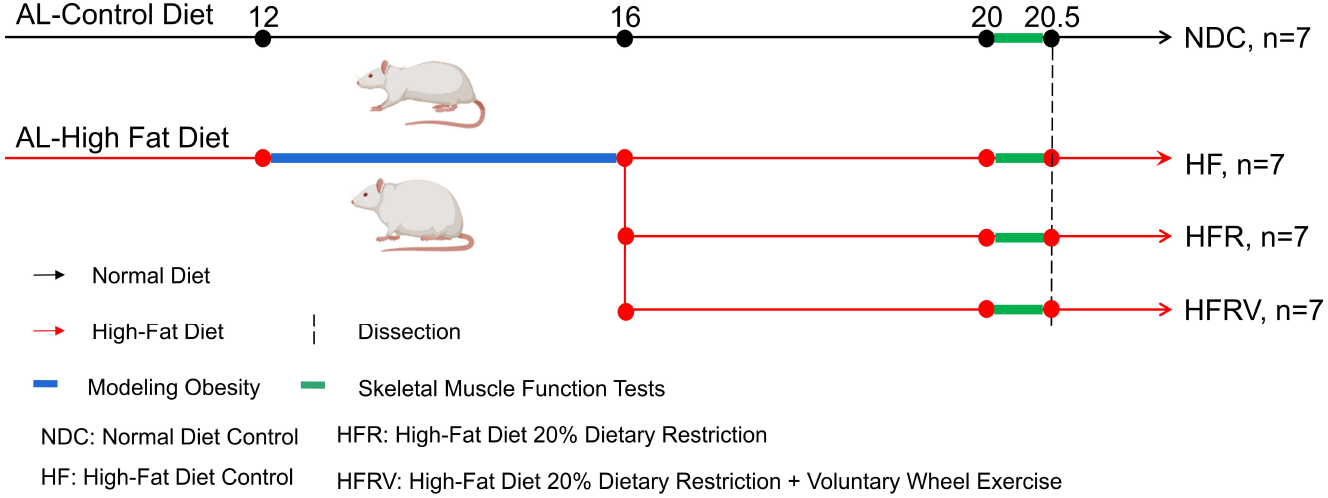
Schematic overview experimental study design.

### 2.2 Body Composition

Mice were anesthetized with 40 mg/kg ketamine and 0.8 mg/kg medetomidine and analyzed for body composition using an InAlyzer Dual-energy X-ray Absorptiometry (DEXA) system (Micro Photonics Inc., PA, USA). The parameters measured included total mass (g), lean mass ratio (%), fat mass (g), lean mass (g), fat content in tissue (%), and fat mass ratio (%).

### 2.3 Skeletal Muscle Function Tests

All mice underwent skeletal muscle function tests before euthanasia. Skeletal muscle function was assessed in four groups of mice using tests for walking speed, endurance, physical activity, and grip strength (Ji et al., 2024).

#### Rota-Rod test

Walking speed was assessed using the Rota-Rod test. An adaptation period was conducted during the first three days, involving a pre-test exercise at 5 rpm for 1 minute, once per day. The official test utilized an acceleration mode, where the speed increased from 5 to 50 rpm over a 5-minute period (Model 76-0770, Harvard Apparatus Inc., Holliston, MA, USA). The latency to fall from the device was recorded. Each mouse was tested three times with a ten-minute interval between tests, and the best performance out of the three trials was used as the outcome measure.

#### Treadmill test

Endurance capacity was evaluated using a treadmill test. Adaptation was performed three days before the official test, consisting of running at 5 cm/s for 5 minutes, once per day, on a 0-degree slope. During the official test, the treadmill speed increased by 1 cm/s every 20 seconds starting from 5 cm/s, with a 0-degree slope. The trial ended when the mouse touched the shock pad (set at 0.6 mA) three times.

#### Voluntary wheel test

Physical activity was measured using the voluntary wheel test. Running distance was recorded using a voluntary wheel (MAN86130, Lafayette Instrument Company, Lafayette, IN, USA), with each wheel rotation corresponding to a distance of 0.4 m. The average running distance over a 5-day period was recorded for each mouse.

#### Inverted-cling grip test

Grip strength was assessed using the inverted-cling grip test. Adaptation was conducted prior to the official test, with the adaptive test performed once a day for three days. During the official test, mice were placed in the center of a wire mesh screen, and a stopwatch was started. The screen was then rotated to an inverted position over 2 seconds, with the mouse’s head descending first, and held 40-50 cm above a padded surface. The time until the mouse fell was recorded. This measurement was repeated three times with ten-minute intervals between tests, and the maximum reading was recorded.

### 2.4 Preservation and Preparation of Skeletal Muscle

The anesthetized mice were euthanized by cervical dislocation. Following euthanasia, the gastrocnemius (GAS) muscle were carefully dissected from the mice. The left GAS was fixed in 10% formalin for 24 hours, then dehydrated and embedded in paraffin. Sections were prepared for Hematoxylin and Eosin (H&E) staining, Prussian blue staining, Masson’s trichrome staining, and immunohistochemistry (IHC). The right GAS was snap-frozen in liquid nitrogen and stored at –80°C for subsequent Western-immunoblot (WB) analyses.

### 2.5 Hematoxylin and Eosin (H&E) staining

Rehydrated tissue sections (5 μm) were stained using ClearView™ hematoxylin (MA0101010, StatLab, McKinney, TX, USA) and ClearView™ eosin (MA0101015, StatLab, McKinney, TX, USA). Subsequently, the sections were dehydrated, cleared, and mounted with neutral resin. Images were acquired using a slide scanner (Axio Scan. Z1, Zeiss) at 200x magnification. For each mouse, a single section of GAS was randomly photographed to obtain 5 images, each covering an area of 0.271mm^2^. The average of these five images was used to represent the cross-sectional area (CSA) of that GAS for the mouse. The CSA was automatically analyzed using “Cellpose,” an open-source deep learning-based segmentation tool (Stringer et al., 2021).

### 2.6 Prussian Blue Staining

Rehydrated tissue sections (5 μm) were stained using Prussian Blue staining kit (G1422, Beijing Solarbio Science & Technology Co., Ltd.). Images were acquired using a slide scanner (Axio Scan. Z1, Zeiss) at 200x magnification. For each mouse, a single section of GAS was randomly photographed to obtain 5 images, each covering an area of 0.271mm^2^. The average of these five images was used to represent the value of that iron accumulation for the mouse. ImageJ software was used for the analysis.

### 2.7 Masson’s Trichrome Staining

Rehydrated tissue sections (5 μm) were stained using Masson’s trichrome staining kit (G1340, Beijing Solarbio Science & Technology Co., Ltd.). Images were acquired using a slide scanner (Axio Scan. Z1, Zeiss) at 200x magnification. For each mouse, a single section of each GAS was randomly photographed to obtain 5 images, each covering an area of 0.271mm^2^. The average of these five images was used to represent the ECM value of that GAS for the mouse. ImageJ software was used for analysis.

### 2.8 Western-Immunoblot (WB)

The red and white GAS were dissected separately (Chen et al., 2015). As previously described (Ji et al., 2024), the GAS was lysed using an EzRIPA Lysis Kit (WSE-7420, ATTO). Protein concentration was determined using the Pierce™ Bicinchoninic Acid Protein Assay Kit (Thermo Scientific, Waltham, MA, USA). Equal amounts of protein samples (40 μg) were separated by sodium dodecyl sulfate-polyacrylamide gel electrophoresis and electro-transferred to nitrocellulose membranes (Bio-Rad Laboratories, Hercules, CA, USA). The membranes were blocked with 5% non-fat milk dissolved in Tris-buffered saline with Tween-20 (TBST; 10 mM Tris, 150 mM NaCl, and 0.1% Tween-20; pH 7.6) for 1.5 hours at room temperature and then incubated with primary antibodies (Table 2) overnight at 4°C. Subsequently, the membranes were incubated with horseradish peroxidase-conjugated secondary antibodies (Table 2) for 1.5 hours at room temperature. The targeted bands were quantified by densitometry using ImageJ software.

**Table 2.** Primary and secondary antibodies.

| Antibodies | Source | Vender | Catalog | Dilution (WB) | Dilution (IHC) |
| --- | --- | --- | --- | --- | --- |
| P53 | Rabbit | Cell signaling | 30313 | 1:500 |  |
| SLC7A11 | Rabbit | Abcam | ab175186 | 1:1000 |  |
| GAPDH | Rabbit | Cell signaling | 5174 | 1:4000 |  |
| GPX4 | Rabbit | Abcam | ab125066 | 1:1000 |  |
| AMPK | Rabbit | Cell signaling | 5831 | 1:500 |  |
| TFR1 | Rabbit | Abcam | ab84036 | 1:500 |  |
| NCOA4 | Rabbit | Invitrogen | PA5-115626 | 1:500 |  |
| Rab7 | Rabbit | Abcam | EPR7589 | 1:1000 |  |
| 4-HNE | Rabbit | Abcam | ab46545 | 1:1000 | 1:200 |
| MDA | Mouse | Abcam | ab243066 | 1:500 | 1:50 |
| Goat anti Rabbit | Goat | Invitrogen | G-21234 | 1:5000 | 1:500 |
| m-IgG Fc | - | SANTA | sc-525409 | 1:5000 | 1:25 |

### 2.9 Immunohistochemistry (IHC)

Following the immunohistochemistry protocol with modifications based on Magaki et al., paraffin-embedded tissue sections (5 μm) were deparaffinized and rehydrated (Magaki et al., 2019). Heat-induced antigen retrieval was performed using 0.01 mol/L sodium citrate buffer (pH 6.0, C1010, Beijing Solarbio Science & Technology Co., Ltd.) in a microwave oven for 20 minutes at 100°C. The sections were then washed twice with Phosphate-buffered saline (PBS; 0.137M NaCl, 0.0027M KCL, 0.010mM Na2HPO4, and 0.002mM KH2PO4; pH 7.2) for 5 minutes each and blocked with 3% hydrogen peroxide (4104-4400, DAE JUNG) for 15 minutes twice to remove endogenous peroxidases. Following three additional 5-minute washes with PBS, the sections were blocked with 100-200 μl of blocking buffer (5% Bovine serum albumin in PBS) for 30 minutes at room temperature. After blocking, the sections were washed three times with PBS for 5 minutes each and incubated with the primary antibody (Table 2) overnight at 4°C. The sections were then washed three times with PBS for 5 minutes each and incubated with the secondary antibody (Table 2) in a humidified chamber for 60 minutes at room temperature. Following three more 5-minute washes with PBS, the sections were incubated with the Pierce™ DAB Substrate Kit (Thermo Scientific, #34002) for 6 minutes and washed three times with PBS for 5 minutes each. Finally, the sections were rehydrated and mounted. Images were acquired using a slide scanner (Axio Scan. Z1, Zeiss) at 200x magnification. For each mouse, a single section of each GAS was randomly photographed to obtain 5 images, each covering an area of 0.271mm^2^. The average of these five images was used to represent the antibody expression value of that GAS for the mouse. ImageJ software was used for analysis.

### 2.10 Statistical Analysis

Statistical analyses were performed using GraphPad Prism software (version 9). Comparisons between two groups were conducted using an unpaired Student’s t-test. For comparisons among three groups, a one-way analysis of variance (ANOVA) followed by a least significant difference (LSD) post-hoc test was employed. To assess the effects of muscle type (white vs. red) and dietary conditions (HF vs. DR vs. DR+Ex), a two-way ANOVA with Bonferroni’s post-hoc test was applied. Outliers were identified and removed using the Z-score approach prior to conducting statistical analyses to ensure the robustness and reliability of the results. All results are expressed as the mean ± standard deviation (SD). Statistical significance was set at p < 0.05, with asterisks indicating the following levels of significance: *p < 0.05, **p < 0.01, ***p < 0.001, and ****p < 0.0001.

## 3. Results

### 3.1 Obesity Evokes Dysfunction and Ferroptosis in Skeletal Muscle

To investigate whether obesity regulates ferroptosis in skeletal muscle, we induced obesity in mice by feeding them a high-fat diet for 14 weeks. All the data are shown in Figure 2.

**Figure 2.**
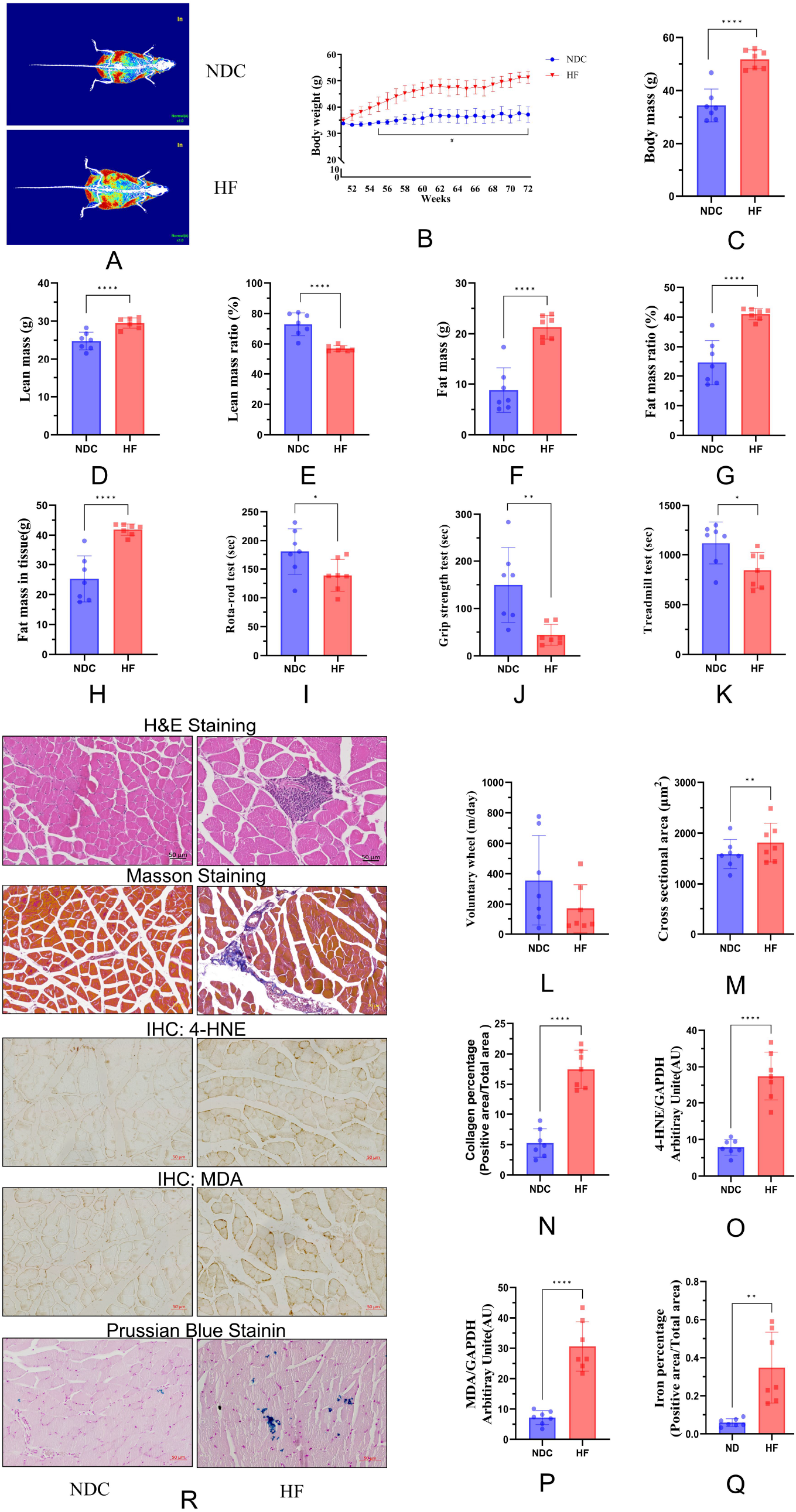
High-fat diet-induced obesity evokes dysfunction and ferroptosis in skeletal muscle. (A) Representative DEXA-scanned images of NDC and HF mice; skeletal muscle, fat tissue, and bone are shown in blue, red, and white, respectively. (B) Weekly body weight changes (group × weeks): NDC vs. HF (*, 54-72 weeks). (C-H) Quantitative data of final total mass, lean mass, lean mass ratio, fat mass, fat mass ratio, and fat mass in tissue, respectively, were acquired by DEXA scanning. (I-L) Skeletal muscle function tests. (I) Walking speed test. (J) Strength test. (K) Endurance test. (L) Physical activity test. (M) Skeletal muscle cross-sectional area in NDC and HF. (N) Skeletal muscle collagen area in NDC and HF. (O) 4-HNE expression levels in skeletal muscle. (P) MDA expression levels in skeletal muscle. (Q) Iron percentage in skeletal muscle. (R) Representative images showing H&E staining, Masson’s trichrome staining, IHC, and Prussian blue staining of GAS. Significant differences are denoted by asterisks: p < 0.05 (*), p < 0.01 (**), p < 0.001 (***), and p < 0.0001 (****). All values are presented as mean ± SD. Normal diet control (NDC, n=7, 20.5-month-old), high-fat diet (HF, n=7, 20.5-month-old). Scale bar = 50 μm.

DEXA analysis revealed that high-fat diet intake led to increases in body weight (P < 0.0001), lean mass (P = 0.0021), fat mass (P < 0.0001), fat mass ratio (P < 0.0001), and fat mass in tissues (P < 0.0001), while the lean mass ratio decreased (P < 0.0001) (Figure 2A-H). These indicators are widely used to evaluate and identify mouse models of obesity (Chen et al., 2012; Koza et al., 2006; Y. Yang et al., 2014; Yeu et al., 2019). Consistent with impaired skeletal muscle function in obese mice (Graham et al., 2019; Vellers et al., 2017), we found that a high-fat diet significantly reduced walking speed (P = 0.0424), grip strength (P = 0.0053), and endurance in mice (P = 0.0217) (Figure 2I-K). These findings further validate the successful establishment of the obese mouse model.

Histological features of ferroptosis include the accumulation of inflammatory cells in tissues (Zhang et al., 2023) and fibrosis (Yu et al., 2020). H&E staining revealed that although the CSA of HF skeletal muscle significantly increased (P = 0.00210), it was accompanied by substantial inflammatory cell infiltration (Figure 2M and R). Additionally, Masson’s trichrome staining indicated a significant increase in skeletal muscle fibrosis in HF compared to NDC (P = 0.0110) (Figure 2N and R). Ferroptosis is highly associated with lipid peroxidation and abnormal iron accumulation (Dixon et al., 2012). As markers of lipid peroxidation, malondialdehyde (MDA) and 4-hydroxynonenal (4-HNE) can reflect the extent of ferroptosis in cells (Chen et al., 2021; Tang et al., 2021). IHC analysis showed elevated levels of lipid peroxidation products 4-HNE (P = 0.0320) and MDA (P = 0.0150) in HF skeletal muscle compared to NDC (Figure 2Q-P and R). Furthermore, Prussian blue staining revealed abnormal iron accumulation in HF skeletal muscle (P = 0.0210) (Figure 2Q and R). These data indicate that obesity, leading to skeletal muscle dysfunction and ferroptosis.

### 3.2 Muscle Type-Specific Regulation of Ferroptosis by GPX4 and NCOA4 in Obesity

Given the observed ferroptosis in skeletal muscle as a result of obesity, we employed WB analysis to investigate the impact of obesity on the expression of representative proteins in 3 ferroptosis-related signaling pathways: amino acid, lipid, and iron metabolism in different muscle types. All the data are shown in Figure 3.

**Figure 4.**
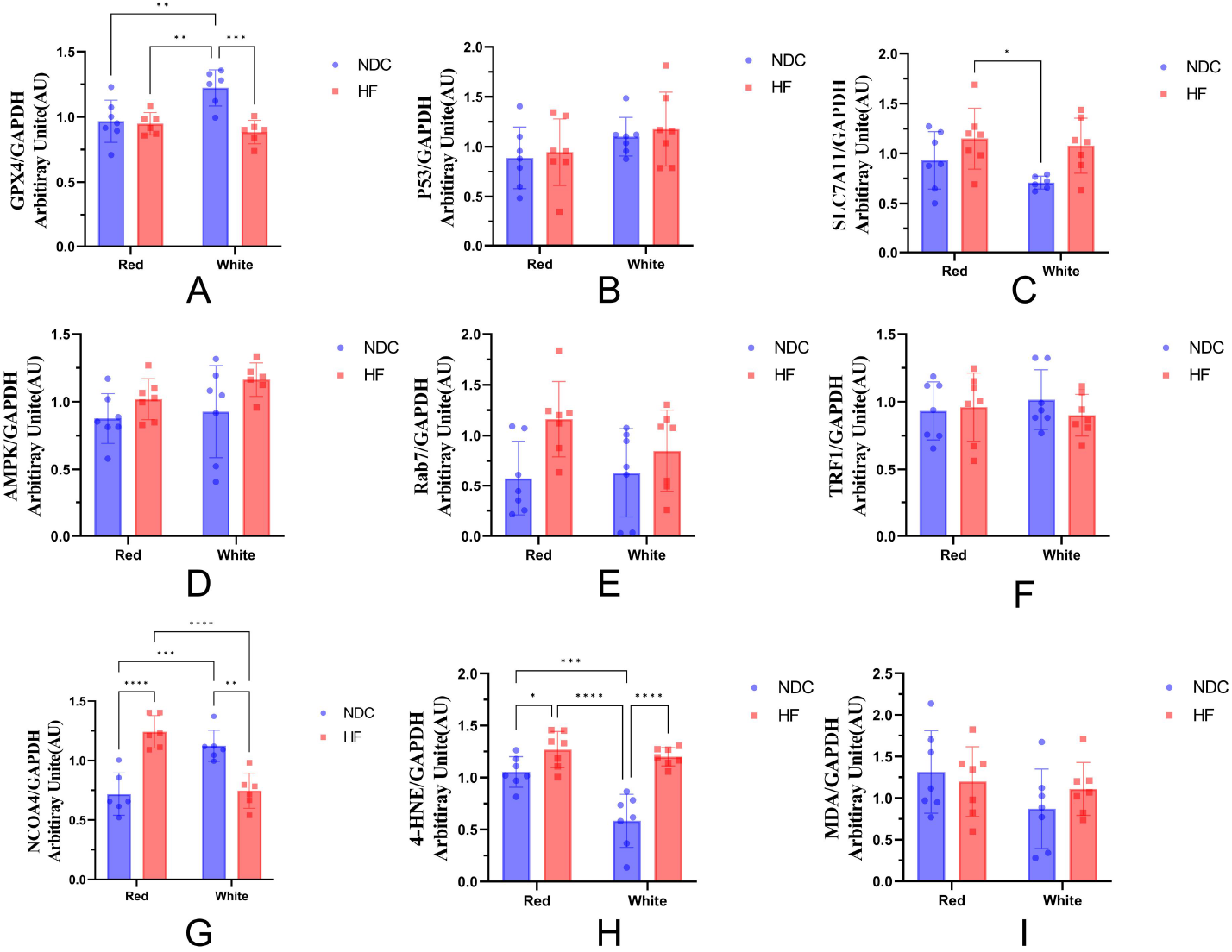
Muscle type-specific regulation of ferroptosis by GPX4 and NOCA4 in high-fat diet-induced obese mice. WB was performed for protein expression levels of ferroptosis signaling pathways, (A): Glutathione peroxidase 4 (GPX4), (B): P53, (C): Solute Carrier Family 7 Member 11 (SLC7A11), (D): AMP-activated protein kinase (AMPK), (E): Ras-related protein Rab-7a (Rab7), (F): Telomeric repeat-binding factor 1 (TRF1), (G) Nuclear Receptor Coactivator 4 (NCOA4), (H): 4-Hydroxynonenal (4-HNE), and (I): Malonaldehyde (MDA). Significant differences are denoted by asterisks: p < 0.05 (*), p < 0.01 (**), p < 0.001 (***) and p < 0.0001 (****). All values are presented as mean ± SD. Normal diet control (NDC, n=7, 20.5-month-old), high-fat diet (HF, n=7, 20.5-month-old).

Under normal diet, the expression of glutathione peroxidase 4 (GPX4) (P = 0.0073) and nuclear receptor coactivator 4 (NCOA4) (P = 0.0007) was significantly higher and 4-HNE (P = 0.002) was lower in white compared to red muscle (Figure 3A and H). Under high-fat diet, the expression of NCOA4 was significantly lower in white compared to red muscle (P < 0.0001) (Figure 3G). Additionally, within the same muscle type, a high-fat diet significantly decreased the expression of GPX4 (P = 0.0007) and NCOA4 (P = 0.0015), and increased 4-HNE (P < 0.0001) in white muscle compared to a normal diet (P < 0.05) (Figure 3A, G, and H). Although a high-fat diet did not alter the expression of GPX4 in red muscle, it significantly increased the expression of NCOA4 (P < 0.0001) and 4-HNE (P = 0.0375) (Figure 3A, G, and H). These data indicated that the regulation of ferroptosis by obesity exhibits skeletal muscle type-specificity. Specifically, obesity regulates ferroptosis by decreasing the GPX4 expression in white muscle and increasing the NCOA4 in red.

### 3.3 DR and DR+Ex Show Improvement and Regulation in Physical Phenotypes and Ferroptosis in Obesity

We identified the regulatory effect of obesity on ferroptosis in skeletal muscle, and subsequently will investigate the intervention mechanisms of DR and DR+Ex on this process. This is because previous studies have widely reported the positive effects of DR and DR+Ex on obesity, lipid peroxidation, and even ferroptosis (Bevilacqua et al., 2005; Chen et al., 2023; Li et al., 2017; Liu et al., 2022; Walsh et al., 2014). All the data are shown in Figure 4.

**Figure 4a.**
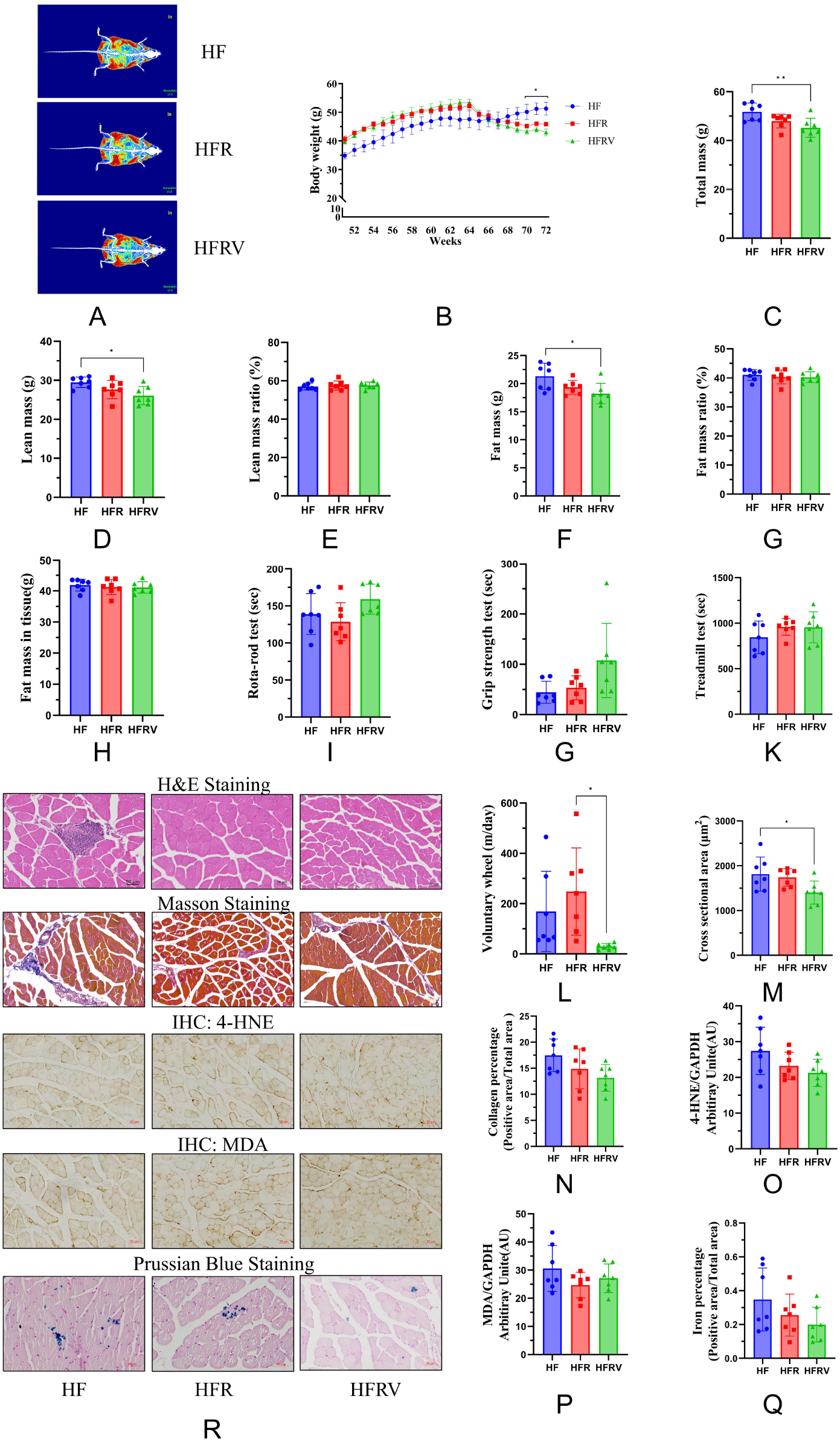
DR and DR+Ex Show Improvement and Regulation in Physical Phenotypes and Ferroptosis in Obesity. A) Representative DEXA-scanned images of NDC and HF mice; skeletal muscle, fat tissue, and bone are shown in blue, red, and white, respectively. (B) Weekly body weight changes (group × weeks): NDC vs. HF (*, 54-72 weeks). (C-H) Quantitative data of final total mass, lean mass, lean mass ratio, fat mass, fat mass ratio, and fat mass in tissue, respectively, were acquired by DEXA scanning. (I-L) Skeletal muscle function tests. (I) Walking speed test. (J) Strength test. (K) Endurance test. (L) Physical activity test. (M) Skeletal muscle cross-sectional area in NDC and HF. (N) Skeletal muscle collagen area in NDC and HF. (O) 4-HNE expression levels in skeletal muscle. (P) MDA expression levels in skeletal muscle. (Q) Iron percentage in skeletal muscle. (R) Representative images showing H&E staining, Masson’s trichrome staining, IHC, and Prussian blue staining of GAS. Significant differences are denoted by asterisks: p < 0.05 (*), p < 0.01 (**), p < 0.001 (***), and p < 0.0001 (****). All values are presented as mean ± SD. High-fat diet (HF, n=7, 20.5-month-old), 20% high-fat dietary restriction (HFR, n=7, 20.5-month-old), 20% high-fat dietary restriction with voluntary wheel running exercise (HFRV, n=7, 20.5-month-old). Scale bar = 50 μm.

**Figure 6.**
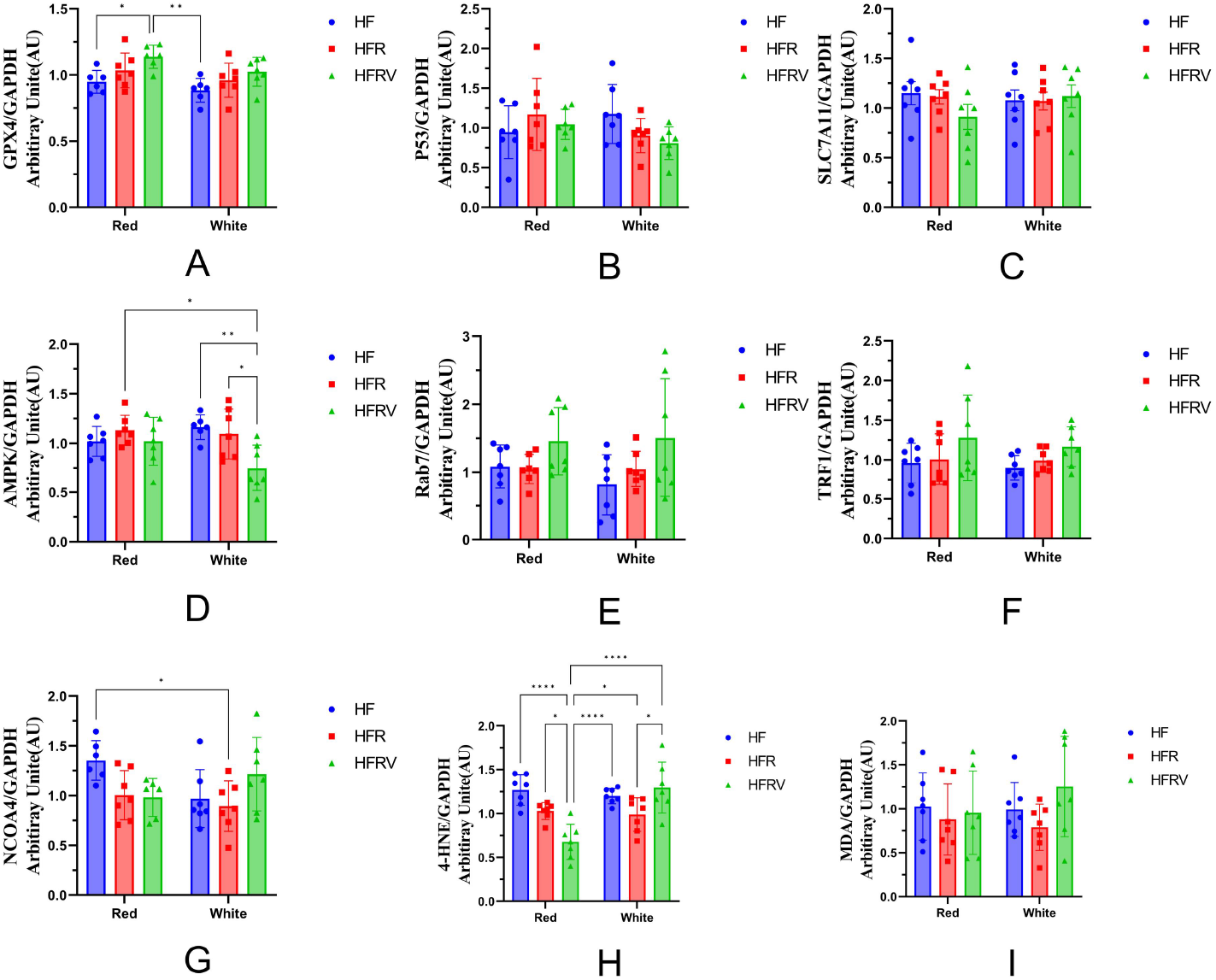
Through CR and CR+Ex muscle type-specific regulation of ferroptosis by GPX4 in high-fat diet-induced obese mice. WB was performed for protein expression levels of ferroptosis signaling pathways, (A): Glutathione peroxidase 4 (GPX4), (B): P53, (C): Solute Carrier Family 7 Member 11 (SLC7A11), (D): AMP-activated protein kinase (AMPK), (E): Ras-related protein Rab-7a (Rab7), (F): Telomeric repeat-binding factor 1 (TRF1), (G) Nuclear Receptor Coactivator 4 (NCOA4), (H): 4-Hydroxynonenal (4-HNE), and (I): Malonaldehyde (MDA). Significant differences are denoted by asterisks: p < 0.05 (*), p < 0.01 (**), p < 0.001 (***) and p < 0.0001 (****). All values are presented as mean ± SD. High-fat diet (HF, n=7), 20% high-fat diet dietary restriction (HFR, n=7), and 20% high-fat diet dietary restriction + voluntary wheel running exercise (HFRV, n=7).

Results showed significant differences in body weight between the HF and HFRV groups at 70 weeks (P = 0.0435) (Figure 4B). Compared to the HF, the HFRV significantly reduced total mass (P = 0.0061), lean mass (P = 0.0171), and fat mass in mice (P = 0.0159), but there were no changes in lean mass ratio, fat mass ratio, or fat mass in tissues (Figure 4C-H). DR and DR+Ex did not appear to effectively improve skeletal muscle function in the HF, HFR, and HFRV groups (Figure 4I-L).

Additionally, H&E staining showed that DR and DR+Ex reduced inflammatory cell infiltration in muscle fibers; however, DR+Ex also decreased the CSA of skeletal muscle (P = 0.0296) (Figure 4M and R). Masson’s trichrome staining and Prussian blue staining indicated fibrosis and iron in skeletal muscle were not significant changes (Figure 4N, Q, and R). Lastly, IHC analysis showed no significant differences in the expression of 4-HNE and MDA among the HF, HFR, and HFRV groups (Figure 4O, P, and R). These data suggest that intervention of DR and DR+Ex has limited effects on improving and regulating physical phenotypes and ferroptosis in obese.

### 3.4 Muscle Type-Specific Regulation of Ferroptosis by GPX4 Through DR and DR+Ex in High-Fat Diet-Induced Obese Mice

Although no significant differences were observed in the histological markers related to ferroptosis, a reduction was noted. Moreover, the detailed of CR and CR+Ex regulatory effects on muscle type-specific ferroptosis are not well understood. Therefore, it is worthwhile to further explore the muscle type-specific regulation of ferroptosis by these interventions through different pathways. The results of the WB analysis are shown in Figure 5.

Under the CR+Ex, the expression of 4-HNE in white muscle was significantly higher compared to red muscle (P < 0.0001) (Figure 5H). In red muscle, the expression of GPX4 was significantly higher in the HFRV group compared to the HF group (P = 0.0471) (Figure 5A); the expression of 4-HNE decreased sequentially in the HF, HFR, and HFRV groups, with significant differences (P < 0.0001 and P = 0.0136) (Figure 5H). In white muscle, the expression of AMP-activated protein kinase (AMPK) was significantly lower in the HFRV group compared to the HF (P = 0.0086) and HFR (P = 0.0126) groups (Figure 5D), and there were significant differences in the expression of 4-HNE between the HFR and HFRV groups (P = 0.0427) (Figure 5H). These data suggest that the regulation of ferroptosis by intervention in high-fat diet-induced obese mice is muscle type-specificity. Specifically, DR+Ex inhibits obesity-induced skeletal muscle ferroptosis by increasing the expression of GPX4 in red muscle.

## 4. Discussion

In recent years, the issue of dysfunction in cells, tissues, and organs caused by obesity has garnered widespread attention. Obesity-induced ferroptosis is considered one of the primary causes of these dysfunctions (Ganz, 2012; Ma et al., 2017; Orr et al., 2014; Zhao et al., 2022). Understanding the regulatory mechanisms of ferroptosis in obesity is significant clinical importance for skeletal muscle-related dysfunction and diseases. This study investigated the impact of obesity on ferroptosis in specific muscle types and the response mechanisms of its signaling pathways to DR and DR+Ex. Our research found that obesity can induce ferroptosis in skeletal muscle, while DR and DR+Ex can regulate this process. More importantly, the regulation of ferroptosis by obesity and its response to intervention are muscle type-specificity. These findings enhance our understanding of the mechanisms of obesity and ferroptosis, providing a scientific basis for further in-depth studies of these diseases and cell death mechanisms.

### 4.1 Appropriateness of an Obesity Mouse Model

Mice at 14-month-old and 20.5-month-old are approximately equivalent to humans aged 30-40 and 60-70 years (Justice et al., 2016). Research has shown that the prevalence of obesity in humans increases significantly after about 35 years of age, reaching a peak around 65 years (Chooi et al., 2019). In our study, mice were fed a high-fat diet starting at 14-month-old and continuing until 20.5-month-old, to simulate sustained obesity in humans beginning at age 35 and persisting until age 65. This experimental design better aligns with the prevalence trends of human obesity, yielding more accurate preclinical data to inform clinical research or the treatment of obesity-induced skeletal muscle-related diseases.

We induced an obese mouse model through 4 months of a high-fat diet (D12451, Research Diets Inc.). This diet has been widely used in the construction of obese mouse models (Armani et al., 2014; Han et al., 2006; Yongbin Yang et al., 2014). Although there is no precise definition for high-fat diet-induced obese mouse models, the vast majority of studies use at least 8 weeks of ad libitum high-fat diet to create obese mouse models (Buettner et al., 2007; Han et al., 2006; Hariri & Thibault, 2010). Additionally, after analyzing the body composition of the mice using DEXA, we found significant increases in body weight, lean mass, fat mass, fat mass in tissues, and fat mass ratio following the high-fat diet. These changes in body composition are consistent with the characteristics of obese mice reported in previous studies (Lin et al., 1977; Mandarim-de-Lacerda et al., 2021; Rogers & Webb, 1980). Moreover, obesity leads to a decline in certain skeletal muscle functions, which is also in line with prior research, indicating that dysfunctional skeletal muscle is a hallmark of obesity (Shortreed et al., 2009; Zhang et al., 2015). Therefore, we conclude that a 4-month high-fat diet can successfully induce obesity, providing an experimental animal model for subsequent study.

### 4.2 Obesity Evokes Ferroptosis in Skeletal Muscle

Recent studies suggest that inflammatory cell infiltration is a hallmark of ferroptosis. Ferroptosis causes lipid peroxidation of cell membranes and cell rupture, and the release of cellular contents (such as high-mobility group protein B1) can activate a local inflammatory response, leading to inflammatory cell infiltration. The study by Dixon et al. pointed out the potential inflammatory role of ferroptosis in various diseases (Dixon et al., 2012). These research findings strongly support that ferroptosis is accompanied by inflammatory cell infiltration. For instance, research by Friedmann Angeli et al. (Friedmann Angeli et al., 2014) demonstrated that ferroptosis leads to acute renal failure, accompanied by significant inflammatory cell infiltration. In the liver, where ferroptosis occurs, inflammatory cell infiltration is also present, as shown in the hemochromatosis mouse model study by Wang et al. (Wang et al., 2017).

Recently, the relationship between ferroptosis and tissue organ fibrosis has been explored in several studies. Ferroptosis, an iron-dependent form of cell death, releases oxidized lipids and cellular contents that can activate fibroblast proliferation, thereby promoting fibrosis. Specifically, cellular contents released during ferroptosis, such as high-mobility group protein B1, can activate fibroblasts and bone marrow-derived fibroblast-like cells. These cells, once activated, secrete large amounts of ECM components like collagen, leading to tissue fibrosis (Kang et al., 2016; Wenzel et al., 2017). Moreover, ferroptosis can activate nuclear factor kappa B (NF-κB) and other pro-inflammatory signaling pathways, promoting the release of pro-fibrotic factors (Dixon et al., 2012; Stockwell et al., 2017). Therefore, ferroptosis is considered a key pathological mechanism in liver and heart fibrosis (Fang et al., 2020; Friedmann Angeli et al., 2014).

Prussian blue staining revealed a significant increase in iron accumulation in the skeletal muscle of obesity. Studies have shown that obese individuals often exhibit lower serum iron levels and elevated hepcidin levels (Ganz, 2012). This increase in hepcidin inhibits iron absorption and release, leading to iron accumulation in the body (Ganz, 2012). Additionally, pro-inflammatory factors secreted in obesity, not only stimulate hepcidin synthesis but also directly affect iron metabolism, increasing iron accumulation in cells and tissues (Hotamisligil, 2006; Weisberg et al., 2003). Ferroptosis is an iron-dependent form of cell death, and its extent can be reflected by the amount of iron accumulation in cells, tissues, and organs. Iron accumulation is a critical trigger for ferroptosis. Excess intracellular iron generates large amounts of ROS through the Fenton Reaction, which induce lipid peroxidation and thereby trigger ferroptosis (Dixon et al., 2012; Stockwell et al., 2017). These research findings strongly support that iron accumulation caused by obesity is one of the factors inducing ferroptosis.

Ferroptosis is accompanied by the accumulation of lipid peroxides and oxidative stress (Dixon et al., 2012). This process is closely related to the expression of 4-HNE and MDA, as both compounds are markers of lipid peroxidation (He et al., 2022; Park et al., 2021). 4-HNE is a highly reactive aldehyde compound produced by lipid peroxidation, primarily derived from the peroxidation of ω-6 polyunsaturated fatty acids (Ayala et al., 2014). It can form adducts with proteins, DNA, and phospholipids, altering their structure and function, thereby causing cellular damage (Murdolo et al., 2023). Similarly, MDA, another product of lipid peroxidation, is commonly used to measure oxidative stress levels. High levels of MDA indicate lipid damage to cell membranes and altered cellular function (Ahmed et al., 2016; Savira et al., 2020). Iron generates highly reactive hydroxyl radicals (•OH) through the Fenton Reaction, which further induce lipid peroxidation, increasing the production of 4-HNE and MDA, enhancing oxidative stress within cells, and promoting ferroptosis (Endale et al., 2023). Thus, the expression levels of 4-HNE and MDA can indirectly reflect ferroptosis activity in tissues and organs.

In summary, obesity leads to inflammation cell infiltration, fibrosis, iron accumulation, and increased lipid peroxides in skeletal muscle. The phenomenon occurrence of these phenomena can confirm that obesity induces ferroptosis in skeletal muscle (Stockwell, 2022).

### 4.3 Regulation of Muscle Type-Specific Ferroptosis Signaling Pathways by Obesity

To better explore the regulatory mechanisms of ferroptosis in skeletal muscle induced by obesity, we used WB to examine the expression of ferroptosis signaling pathways in different types of muscle.

GPX4 plays a key regulatory role in ferroptosis through amino acid metabolism by inhibiting the cell death process via the reduction of lipid peroxides accumulation (Stockwell et al., 2017). The deficiency or inactivation of GPX4 leads to an increase in lipid peroxides, thereby triggering ferroptosis (W. S. Yang et al., 2014). In obesity, the expression of GPX4 in white muscle significantly decreases, while its expression in red muscle remains unaffected. This may be due to the higher sensitivity of white muscle to lipid peroxidation. A high-fat diet increases oxidative stress in white muscle, leading to GPX4 depletion, whereas red muscle, with its stronger antioxidant capacity, can maintain GPX4 levels (Katunga et al., 2015; Zhuang et al., 2021). Additionally, higher metabolic activity and lipid accumulation in white muscle may also contribute to the significant decrease in GPX4 expression (Shi et al., 2024).

NCOA4 mediates ferritinophagy, the process of degrading ferritin to release stored iron, thereby increasing intracellular free iron content. This promotes iron-catalyzed Fenton Reactions, leading to lipid peroxidation and ferroptosis (Dowdle et al., 2014; Mancias et al., 2014). Studies have shown that the absence of NCOA4 significantly reduces the occurrence of ferroptosis, further demonstrating its crucial role in the ferroptosis process (Dowdle et al., 2014; Mancias et al., 2014). Obesity impacts the expression of NCOA4 differently in red and white muscles, with an increase in red muscle and a significant decrease in white muscle. White muscle primarily relies on anaerobic metabolism and is more sensitive to lipid peroxidation and metabolic stress. The increased metabolic stress caused by a high-fat diet enhances the white muscle cells’ need for oxidative stress defense, possibly downregulating NCOA4 expression to reduce ferritinophagy activities, thereby protecting the cells damage from further ferroptosis. This hypothesis requires further in-depth research.

Obesity leads to increased lipid accumulation and oxidative stress in both red and white muscles, significantly enhancing the production of 4-HNE (Hart et al., 2018; Shi et al., 2024). Obesity also weakens antioxidant defense mechanisms, such as reducing the activity of glutathione and superoxide dismutase, further exacerbating lipid peroxidation (Hart et al., 2018). It is of paramount importance to note that the skeletal muscle type specific expression of GPX4 and NCOA4 as upstream and regulatory signals of 4-HNE was significantly modulated by high-fat-induced obesity. Consequently, we propose that the upregulation of 4-HNE in red and white muscle due to high-fat induced obesity is caused by significant alterations in the expression of NCOA4 and GPX4, separately.

In obesity, the expression of 4-HNE significantly increased in both red and white muscles, whereas the expression of MDA did not show significant changes. This discrepancy may be due to the differences in the generation and metabolism processes of 4-HNE and MDA. The production of 4-HNE is highly influenced by ROS. Obesity increases lipid peroxidation reactions and ROS production, thereby significantly elevating 4-HNE levels (Morris et al., 2008; Ramana et al., 2014). Additionally, 4-HNE is highly reactive and can form stable adducts with various biomolecules such as proteins, DNA, and phospholipids, leading to its accumulation in tissues (Ayala et al., 2014; Morris et al., 2008). However, the production of MDA is relatively stable, and its metabolic pathways are more diverse and efficient. For example, MDA can be metabolized through various pathways, such as being converted into water-soluble products by glutathione-S-transferase for excretion or being transformed into less reactive compounds (Esterbauer et al., 1991). These metabolic pathways maybe result in insignificant changes in MDA levels under obesity (Del Rio et al., 2005).

Additionally, experimental techniques can affect the expression of MDA. When using IHC and WB to detect MDA, IHC showed a significant increase in MDA after a high-fat diet, whereas WB did not show the same results. This discrepancy may arise from differences in the sensitivity and specificity of the detection methods (Del Rio et al., 2005). IHC can detect local accumulation of MDA in specific cell types or tissue regions, while WB measures overall protein levels, which may dilute signals of local high expression. Furthermore, differences in sample processing and extraction methods may lead to dilution or degradation of locally high concentrations of MDA in WB, affecting the detection results. These factors collectively explain why IHC can show a significant increase in MDA after a high-fat diet, while WB does not detect the same changes.

### 4.4 Regulation of Ferroptosis by DR and DR+Ex

We found that after 8 weeks of intervention, although some physical phenotypes and ferroptosis-related characteristics of the mice improved, there was no significant difference. This may be related to the intensity of the intervention and the metabolism of mice. Higher-intensity caloric restriction can produce more pronounced positive effects, but it is not suitable for aged mice (Schädel et al., 2023). Using treadmills and other experimental equipment can force mice to exercise and achieve greater exercise benefits, but this can impose stress on the mice and even lead to mortality (Narath et al., 2001). Additionally, aging causes mice to become frailer, and obesity exacerbates this negative effect, further increasing their mortality rate (Zhang et al., 2015). Therefore, we chose a low-intensity CR combined with or without voluntary wheel running intervention for the mice to ensure the survival rate of the mice and the successful completion of the experiment. Furthermore, aging (Rattan & Derventzi, 1991) and obesity (de Wilde et al., 2009) lower the body’s metabolic rate and responsiveness to external stimuli. The combined effects of these factors are the primary reasons for the lack of significant intervention results.

### 4.5 DR+Ex Regulated Ferroptosis Expression in Red Muscle

Although no significant regulatory effects of DR and DR+Ex on ferroptosis signals (4-HNE and MDA) in the wholly skeletal muscles of obese mice were found, expression did reduced. Therefore, we used WB analysis to examine the regulatory mechanisms of DR and DR+Ex on ferroptosis signals in different muscle types. The results showed that after DR+Ex, the expression of GPX4 in red muscle significantly increased in obese mice, whereas it did not in white muscle. This may be related to the oxidative stress, metabolic demands, and specific gene regulatory mechanisms of different skeletal muscle types.

Firstly, red muscle primarily relies on aerobic metabolism and is rich in mitochondria, adapting to prolonged low-intensity exercise. Studies have shown that antioxidant mechanisms in red muscle are more prominent, potentially leading to the easier expression of GPX4 in response to oxidative stress in this environment (Rinnankoski-Tuikka et al., 2012). Secondly, DR and Ex lead to increased aerobic metabolism activity in skeletal muscle, requiring more antioxidant protection to prevent lipid peroxidation damage (Civitarese et al., 2007; Galassetti et al., 2006). As a key antioxidant enzyme, GPX4 is upregulated in this context, while white muscle, which primarily relies on glycolysis for energy, has lower oxidative stress, and its GPX4 expression maybe does not significantly increase. Lastly, DR and Ex can regulate gene expression through specific signaling pathways such as AMPK and PGC-1α. These pathways are more active in red muscle (Gouspillou et al., 2014; Jørgensen et al., 2007; Pengam et al., 2020) and are related to the expression of antioxidant enzymes. In summary, the increased expression of GPX4 in red muscle may be due to its high sensitivity to oxidative stress, higher metabolic demands, and specific gene regulation induced by endurance exercise. In contrast, white muscle, with its different metabolic characteristics, did not show the same changes in GPX4 expression.

We found that after DR+Ex in obese mice, the expression of AMPK decreased in white muscle but not in red muscle. This may be related to the metabolic differences between muscle fiber types, exercise-induced specific regulation, and the different impacts of diet and exercise on metabolic regulation. Red muscle primarily relies on aerobic metabolism, is rich in mitochondria, and is adapted to prolonged low-intensity exercise, maintaining high AMPK activity to meet its metabolic demands and antioxidant capacity. Under conditions of exercise and dietary restriction, AMPK expression remains stable in red muscle. In contrast, white muscle primarily relies on glycolysis for energy, has lower oxidative stress at resting stat, and therefore requires less AMPK, after intervention leading to a possible decrease in its expression (Jørgensen et al., 2007).

After DR and DR+Ex in obese mice, the expression of 4-HNE decreased in red muscle but not in white muscle. This difference is related to the metabolic differences between skeletal muscle types, specific gene regulation, and the specific effects of diet and exercise on red muscle. Red muscle primarily relies on aerobic metabolism, is rich in mitochondria, and is adapted to prolonged low-intensity exercise. Consequently, its metabolic activity and oxidative stress are higher, necessitating stronger antioxidant protection mechanisms. Studies have shown that red muscle significantly increases antioxidant enzyme activity under DR and Ex, reducing the production of oxidative stress markers such as 4-HNE (Jørgensen et al., 2007). Moreover, exercise regulates gene expression through specific signaling pathways which are more active in red muscle. This promotes the expression and activity of antioxidant enzymes, further reducing levels of oxidative stress markers (Sriwijitkamol et al., 2006).

We found that as an upstream signal and regulatory molecule of 4-HEN, the expression of GPX4 in red muscle significantly increased after intervention. Therefore, we suggest that DR and DR+Ex can regulate ferroptosis by altering the expression of GPX4 in red muscle, which in turn modulates the downstream expression of 4-HEN. However, AMPK, also an upstream regulatory molecule of 4-HEN, showed inconsistent expression changes with 4-HNE in white muscle. After a high-fat diet, the regulatory effect of AMPK on 4-HNE in white muscle was impaired, likely due to metabolic disturbances caused by obesity. This warrants further investigation in future studies.

## 5. Conclusion

Obesity induced by a high-fat diet leads to ferroptosis in the skeletal muscle of mice. The mechanism is muscle type-specific, with NCOA4 expression upregulated in red muscle and GPX4 expression upregulated in white muscle, ultimately triggering ferroptosis. Additionally, DR and DR+Ex can partially mitigate ferroptosis in the skeletal muscle of obese mice induced by a high-fat diet. This mechanism is achieved by regulating the expression of GPX4 in red muscle. Overall, we have identified muscle type-specific regulatory mechanisms of ferroptosis, providing molecular biology-related scientific data for the prevention and treatment of related diseases in the future.

## 6. limitation

The study focuses on the gastrocnemius muscle, which may not fully represent the effects of dietary restriction and exercise on other muscle groups, as different muscles may respond differently to these interventions. Additionally, the duration of dietary restriction and exercise intervention was limited to eight weeks. Longer-term studies are necessary to understand the chronic effects and sustainability of these interventions on ferroptosis and overall muscle health. Furthermore, incorporating advanced experimental tools, such as single-cell sequencing, could provide a more detailed and microscopic analysis of the results.

## 7. WB representative picture

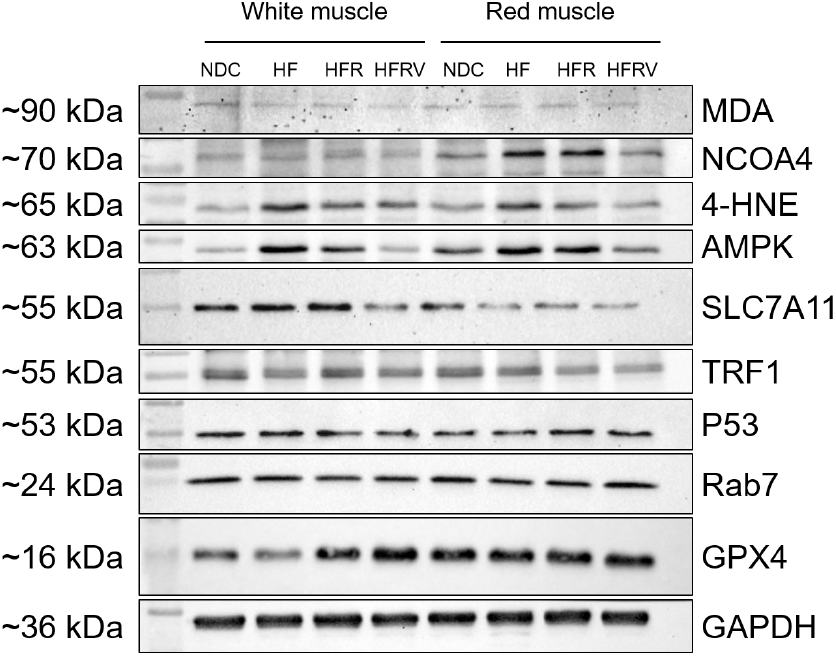

## 8. Ethical approval and consent to participate

All the procedures followed in this experiment were approved by the Institutional Animal Care and Use Committee (IACUC) of Hanyang University (HYU 2021-0066A).

## 9. Consent for publication

Not applicable.

## 10. Availability of data and materials

The data used to support the findings of this study are presented here. Any further data requirements are available from the corresponding author upon request.

## 13. Author’ s contributions

F.J.J. and J.H.K. conceived and designed the study. F.J.J., Y.J.P. and H.s.L. performed the experiments. F.J.J. and H.s.L. analyzed the data and prepared the figures. F.J.J. and J.H.K. interpreted the results, drafted, edited, and revised the manuscript. J.H.K. acquired funding. All authors approved the final version of the manuscript.

## 14. Acknowledgments

We thank Jian Guo (Hanyang University, Korea) for his generous help in dissecting the mice.

We are grateful to Prof. Dr. Gwang-woong Go (Hanyang University, Korea) for generously providing the DEXA machine for body composition analysis of the mice.

We thank the support of Skill Learning from Kaixin Doctor and MASCU (Medical Association with Science, Creativity, and Unity), Inc, Shenzhen, China.

## Reference

Ahmed, S., Maher, F., & Naji, N. (2016). Effect of leptin and oxidative stress in the blood of obese individuals. Biochem Anal Biochem, 5(288), 2161–1009.

Armani, A., Cinti, F., Marzolla, V., Morgan, J., Cranston, G. A., Antelmi, A., Carpinelli, G., Canese, R., Pagotto, U., & Quarta, C. (2014). Mineralocorticoid receptor antagonism induces browning of white adipose tissue through impairment of autophagy and prevents adipocyte dysfunction in high-fat-diet-fed mice. The FASEB Journal, 28(8), 3745–3757.

Ayala, A., Muñoz, M. F., & Argüelles, S. (2014). Lipid peroxidation: production, metabolism, and signaling mechanisms of malondialdehyde and 4-hydroxy-2-nonenal. Oxidative Medicine and Cellular Longevity, 2014.

Bevilacqua, L., Ramsey, J. J., Hagopian, K., Weindruch, R., & Harper, M.-E. (2005). Long-term caloric restriction increases UCP3 content but decreases proton leak and reactive oxygen species production in rat skeletal muscle mitochondria. American Journal of Physiology-Endocrinology and Metabolism, 289(3), E429–E438.

Blüher, M. (2019). Obesity: global epidemiology and pathogenesis. Nature Reviews Endocrinology, 15(5), 288–298.

Buettner, R., Schölmerich, J., & Bollheimer, L. C. (2007). High-fat diets: modeling the metabolic disorders of human obesity in rodents. Obesity, 15(4), 798–808.

Chen, D., Wang, Y., & Chin, E. R. (2015). Activation of the endoplasmic reticulum stress response in skeletal muscle of G93A*SOD1 amyotrophic lateral sclerosis mice. Front Cell Neurosci, 9, 170. 10.3389/fncel.2015.00170

Chen, J., Zhu, T., Yu, D., Yan, B., Zhang, Y., Jin, J., Yang, Z., Zhang, B., Hao, X., & Chen, Z. (2023). Moderate intensity of treadmill exercise rescues TBI-induced ferroptosis, neurodegeneration, and cognitive impairments via suppressing STING pathway. Molecular Neurobiology, 60(9), 4872–4896.

Chen, W., Wilson, J. L., Khaksari, M., Cowley, M. A., & Enriori, P. J. (2012). Abdominal fat analyzed by DEXA scan reflects visceral body fat and improves the phenotype description and the assessment of metabolic risk in mice. Am J Physiol Endocrinol Metab, 303(5), E635–643. 10.1152/ajpendo.00078.2012

Chen, X., Comish, P. B., Tang, D., & Kang, R. (2021). Characteristics and Biomarkers of Ferroptosis. Front Cell Dev Biol, 9, 637162. 10.3389/fcell.2021.637162

Chooi, Y. C., Ding, C., & Magkos, F. (2019). The epidemiology of obesity. Metabolism, 92, 6–10. 10.1016/j.metabol.2018.09.005

Civitarese, A. E., Carling, S., Heilbronn, L. K., Hulver, M. H., Ukropcova, B., Deutsch, W. A., Smith, S. R., & Ravussin, E. (2007). Calorie restriction increases muscle mitochondrial biogenesis in healthy humans. PLoS medicine, 4(3), e76.

Collins, K. H., Herzog, W., MacDonald, G. Z., Reimer, R. A., Rios, J. L., Smith, I. C., Zernicke, R. F., & Hart, D. A. (2018). Obesity, metabolic syndrome, and musculoskeletal disease: common inflammatory pathways suggest a central role for loss of muscle integrity. Frontiers in physiology, 9, 112.

de Wilde, J., Smit, E., Mohren, R., Boekschoten, M. V., de Groot, P., van den Berg, S. A., Bijland, S., Voshol, P. J., van Dijk, K. W., de Wit, N. W., Bunschoten, A., Schaart, G., Hulshof, M. F., & Mariman, E. C. (2009). An 8-week high-fat diet induces obesity and insulin resistance with small changes in the muscle transcriptome of C57BL/6J mice. J Nutrigenet Nutrigenomics, 2(6), 280–291. 10.1159/000308466

Del Rio, D., Stewart, A. J., & Pellegrini, N. (2005). A review of recent studies on malondialdehyde as toxic molecule and biological marker of oxidative stress. Nutr Metab Cardiovasc Dis, 15(4), 316–328. 10.1016/j.numecd.2005.05.003

Ding, H., Chen, S., Pan, X., Dai, X., Pan, G., Li, Z., Mai, X., Tian, Y., Zhang, S., Liu, B., Cao, G., Yao, Z., Yao, X., Gao, L., Yang, L., Chen, X., Sun, J., Chen, H., Han, M., … Xie, L. (2021). Transferrin receptor 1 ablation in satellite cells impedes skeletal muscle regeneration through activation of ferroptosis. J Cachexia Sarcopenia Muscle, 12(3), 746–768. 10.1002/jcsm.12700

Dixon, S. J., Lemberg, K. M., Lamprecht, M. R., Skouta, R., Zaitsev, E. M., Gleason, C. E., Patel, D. N., Bauer, A. J., Cantley, A. M., & Yang, W. S. (2012). Ferroptosis: an iron-dependent form of nonapoptotic cell death. cell, 149(5), 1060–1072.

Dowdle, W. E., Nyfeler, B., Nagel, J., Elling, R. A., Liu, S., Triantafellow, E., Menon, S., Wang, Z., Honda, A., Pardee, G., Cantwell, J., Luu, C., Cornella-Taracido, I., Harrington, E., Fekkes, P., Lei, H., Fang, Q., Digan, M. E., Burdick, D., Murphy, L. O. (2014). Selective VPS34 inhibitor blocks autophagy and uncovers a role for NCOA4 in ferritin degradation and iron homeostasis in vivo. Nature cell biology, 16(11), 1069–1079. 10.1038/ncb3053

Endale, H. T., Tesfaye, W., & Mengstie, T. A. (2023). ROS induced lipid peroxidation and their role in ferroptosis. Frontiers in cell and Developmental Biology, 11.

Esterbauer, H., Schaur, R. J., & Zollner, H. (1991). Chemistry and biochemistry of 4-hydroxynonenal, malonaldehyde and related aldehydes. Free Radic Biol Med, 11(1), 81–128. 10.1016/0891-5849(91)90192-6

Fang, D., Wang, Y., Zhang, Z., Yang, D., Gu, D., He, B., Zhang, X., He, D., Wang, H., & Jose, P. A. (2021). Calorie restriction protects against contrast-induced nephropathy via SIRT1/GPX4 activation. Oxidative Medicine and Cellular Longevity, 2021(1), 2999296.

Fang, X., Cai, Z., Wang, H., Han, D., Cheng, Q., Zhang, P., Gao, F., Yu, Y., Song, Z., & Wu, Q. (2020). Loss of cardiac ferritin H facilitates cardiomyopathy via Slc7a11-mediated ferroptosis. Circulation research, 127(4), 486–501.

Franco-Romero, A., Sandri, M., & Schiaffino, S. (2024). Autophagy in Skeletal Muscle. Cold Spring Harbor Perspectives in Biology, a041565.

Friedmann Angeli, J. P., Schneider, M., Proneth, B., Tyurina, Y. Y., Tyurin, V. A., Hammond, V. J., Herbach, N., Aichler, M., Walch, A., & Eggenhofer, E. (2014). Inactivation of the ferroptosis regulator Gpx4 triggers acute renal failure in mice. Nature cell biology, 16(12), 1180–1191.

Galassetti, P. R., Nemet, D., Pescatello, A., Rose-Gottron, C., Larson, J., & Cooper, D. M. (2006). Exercise, caloric restriction, and systemic oxidative stress. Journal of investigative medicine, 54(2), 67–75.

Ganz, T. (2012). Macrophages and systemic iron homeostasis. Journal of innate immunity, 4(5-6), 446–453.

Gouspillou, G., Sgarioto, N., Norris, B., Barbat-Artigas, S., Aubertin-Leheudre, M., Morais, J. A., Burelle, Y., Taivassalo, T., & Hepple, R. T. (2014). The relationship between muscle fiber type-specific PGC-1α content and mitochondrial content varies between rodent models and humans. PloS one, 9(8), e103044. 10.1371/journal.pone.0103044

Graham, L. C., Grabowska, W. A., Chun, Y., Risacher, S. L., Philip, V. M., Saykin, A. J., Rizzo, S. J. S., Howell, G. R., & Initiative, A. s. D. N. (2019). Exercise prevents obesity-induced cognitive decline and white matter damage in mice. Neurobiology of aging, 80, 154–172.

Guerrero-Hue, M., García-Caballero, C., Palomino-Antolín, A., Rubio-Navarro, A., Vázquez-Carballo, C., Herencia, C., Martín-Sanchez, D., Farré-Alins, V., Egea, J., Cannata, P., Praga, M., Ortiz, A., Egido, J., Sanz, A. B., & Moreno, J. A. (2019). Curcumin reduces renal damage associated with rhabdomyolysis by decreasing ferroptosis-mediated cell death. Faseb j, 33(8), 8961–8975. 10.1096/fj.201900077R

Han, C., Lee, S., Lim, S., Kong, H., Kim, S., Lee, S., & Chang, J. (2006). Studies on production of high fat diet induced obesity C57BL/6NCrjBgi mice. Laboratory Animal Research, 22(3), 221–226.

Hariri, N., & Thibault, L. (2010). High-fat diet-induced obesity in animal models. Nutrition research reviews, 23(2), 270–299.

Hart, C. R., Layec, G., Trinity, J. D., Kwon, O. S., Zhao, J., Reese, V. R., Gifford, J. R., & Richardson, R. S. (2018). Increased skeletal muscle mitochondrial free radical production in peripheral arterial disease despite preserved mitochondrial respiratory capacity. Exp Physiol, 103(6), 838–850. 10.1113/ep086905

He, F., Huang, X., Wei, G., Lin, X., Zhang, W., Zhuang, W., He, W., Zhan, T., Hu, H., & Yang, H. (2022). Regulation of ACSL4-catalyzed lipid peroxidation process resists cisplatin ototoxicity. Oxidative Medicine and Cellular Longevity, 2022.

He, L. P., Zhou, Z. X., & Li, C. P. (2023). Narrative review of ferroptosis in obesity. Journal of Cellular and Molecular Medicine, 27(7), 920–926.

Hotamisligil, G. S. (2006). Inflammation and metabolic disorders. Nature, 444(7121), 860–867.

Huang, Y., Wu, B., Shen, D., Chen, J., Yu, Z., & Chen, C. (2021). Ferroptosis in a sarcopenia model of senescence accelerated mouse prone 8 (SAMP8). Int J Biol Sci, 17(1), 151–162. 10.7150/ijbs.53126

Ji, F., Park, J. H., Rheem, H., & Kim, J.-H. (2024). Overlapping and Distinct Physical and Biological Phenotypes Related to Pure Frailty and Frail Obesity. Bioscience Reports, BSR20240784.

Jørgensen, S. B., Treebak, J. T., Viollet, B., Schjerling, P., Vaulont, S., Wojtaszewski, J. F., & Richter, E. A. (2007). Role of AMPKα2 in basal, training-, and AICAR-induced GLUT4, hexokinase II, and mitochondrial protein expression in mouse muscle. American Journal of Physiology-Endocrinology and Metabolism, 292(1), E331–E339.

Justice, J. N., Cesari, M., Seals, D. R., Shively, C. A., & Carter, C. S. (2016). Comparative Approaches to Understanding the Relation Between Aging and Physical Function. J Gerontol A Biol Sci Med Sci, 71(10), 1243–1253. 10.1093/gerona/glv035

Kalinkovich, A., & Livshits, G. (2017). Sarcopenic obesity or obese sarcopenia: a cross talk between age-associated adipose tissue and skeletal muscle inflammation as a main mechanism of the pathogenesis. Ageing research reviews, 35, 200–221.

Kang, R., Zeng, L., Xie, Y., Yan, Z., Zhou, B., Cao, L., Klionsky, D. J., Tracey, K. J., Li, J., & Wang, H. (2016). A novel PINK1-and PARK2-dependent protective neuroimmune pathway in lethal sepsis. Autophagy, 12(12), 2374–2385.

Katunga, L. A., Gudimella, P., Efird, J. T., Abernathy, S., Mattox, T. A., Beatty, C., Darden, T. M., Thayne, K. A., Alwair, H., Kypson, A. P., Virag, J. A., & Anderson, E. J. (2015). Obesity in a model of gpx4 haploinsufficiency uncovers a causal role for lipid-derived aldehydes in human metabolic disease and cardiomyopathy. Mol Metab, 4(6), 493–506. 10.1016/j.molmet.2015.04.001

Khachatoorian, Y., & Samara, A. (2018). Differential effects of dietary restriction combined with exercise vs dietary restriction alone on visceral and subcutaneous adipose tissues: A systematic review. Obesity Medicine, 9, 7–17.

Kim, J. D., McCarter, R. J., & Yu, B. P. (1996). Influence of age, exercise, and dietary restriction on oxidative stress in rats. Aging (Milano), 8(2), 123–129. 10.1007/bf03339566

Koza, R. A., Nikonova, L., Hogan, J., Rim, J. S., Mendoza, T., Faulk, C., Skaf, J., & Kozak, L. P. (2006). Changes in gene expression foreshadow diet-induced obesity in genetically identical mice. PLoS Genet, 2(5), e81. 10.1371/journal.pgen.0020081

Kvedaras, M., Minderis, P., Krusnauskas, R., & Ratkevicius, A. (2020). Effects of ten-week 30% caloric restriction on metabolic health and skeletal muscles of adult and old C57BL/6J mice. Mechanisms of Ageing and Development, 190, 111320.

Li, C., Feng, F., Xiong, X., Li, R., & Chen, N. (2017). Exercise coupled with dietary restriction reduces oxidative stress in male adolescents with obesity. Journal of sports sciences, 35(7), 663–668.

Lin, P.-Y., Romsos, D. R., & Leveille, G. A. (1977). Food intake, body weight gain, and body composition of the young obese (ob/ob) mouse. The Journal of Nutrition, 107(9), 1715–1723.

Liu, T., Cui, Y., Dong, S., Kong, X., Xu, X., Wang, Y., Wan, Q., & Wang, Q. (2022). Treadmill training reduces cerebral ischemia-reperfusion injury by inhibiting ferroptosis through activation of SLC7A11/GPX4. Oxidative Medicine and Cellular Longevity, 2022.

Ma, X., Pham, V. T., Mori, H., MacDougald, O. A., Shah, Y. M., & Bodary, P. F. (2017). Iron elevation and adipose tissue remodeling in the epididymal depot of a mouse model of polygenic obesity. PloS one, 12(6), e0179889.

Magaki, S., Hojat, S. A., Wei, B., So, A., & Yong, W. H. (2019). An Introduction to the Performance of Immunohistochemistry. Methods Mol Biol, 1897, 289–298. 10.1007/978-1-4939-8935-5_25

Mancias, J. D., Wang, X., Gygi, S. P., Harper, J. W., & Kimmelman, A. C. (2014). Quantitative proteomics identifies NCOA4 as the cargo receptor mediating ferritinophagy. Nature, 509(7498), 105–109.

Mandarim-de-Lacerda, C. A., Del Sol, M., Vazquez, B., & Aguila, M. B. (2021). Mice as an animal model for the study of adipose tissue and obesity. Int J Morphol, 39(6), 1521–1528.

Mizushima, N. (2007). Autophagy: process and function. Genes & development, 21(22), 2861–2873.

Morris, R. T., Laye, M. J., Lees, S. J., Rector, R. S., Thyfault, J. P., & Booth, F. W. (2008). Exercise-induced attenuation of obesity, hyperinsulinemia, and skeletal muscle lipid peroxidation in the OLETF rat. J Appl Physiol (1985), 104(3), 708–715. 10.1152/japplphysiol.01034.2007

Murdolo, G., Bartolini, D., Tortoioli, C., Vermigli, C., Piroddi, M., & Galli, F. (2023). Accumulation of 4-Hydroxynonenal Characterizes Diabetic Fat and Modulates Adipogenic Differentiation of Adipose Precursor Cells. International Journal of Molecular Sciences, 24(23), 16645.

Narath, E., Skalicky, M., & Viidik, A. (2001). Voluntary and forced exercise influence the survival and body composition of ageing male rats differently. Experimental gerontology, 36(10), 1699–1711.

Orr, J. S., Kennedy, A., Anderson-Baucum, E. K., Webb, C. D., Fordahl, S. C., Erikson, K. M., Zhang, Y., Etzerodt, A., Moestrup, S. K., & Hasty, A. H. (2014). Obesity alters adipose tissue macrophage iron content and tissue iron distribution. Diabetes, 63(2), 421–432.

Ostler, J. E., Maurya, S. K., Dials, J., Roof, S. R., Devor, S. T., Ziolo, M. T., & Periasamy, M. (2014). Effects of insulin resistance on skeletal muscle growth and exercise capacity in type 2 diabetic mouse models. American Journal of Physiology-Endocrinology and Metabolism, 306(6), E592–E605.

Park, M. W., Cha, H. W., Kim, J., Kim, J. H., Yang, H., Yoon, S., Boonpraman, N., Yi, S. S., Yoo, I. D., & Moon, J.-S. (2021). NOX4 promotes ferroptosis of astrocytes by oxidative stress-induced lipid peroxidation via the impairment of mitochondrial metabolism in Alzheimer’s diseases. Redox biology, 41, 101947.

Pengam, M., Moisan, C., Simon, B., Guernec, A., Inizan, M., & Amérand, A. (2020). Training protocols differently affect AMPK-PGC-1α signaling pathway and redox state in trout muscle. Comp Biochem Physiol A Mol Integr Physiol, 243, 110673. 10.1016/j.cbpa.2020.110673

Ramana, K. V., Srivastava, S., & Singhal, S. S. (2014). Lipid peroxidation products in human health and disease 2014. Oxid Med Cell Longev, 2014, 162414. 10.1155/2014/162414

Rattan, S. I., & Derventzi, A. (1991). Altered cellular responsiveness during ageing. Bioessays, 13(11), 601–606.

Rinnankoski-Tuikka, R., Silvennoinen, M., Torvinen, S., Hulmi, J. J., Lehti, M., Kivelä, R., Reunanen, H., & Kainulainen, H. (2012). Effects of high-fat diet and physical activity on pyruvate dehydrogenase kinase-4 in mouse skeletal muscle. Nutrition & metabolism, 9, 1–13.

Rogers, P., & Webb, G. P. (1980). Estimation of body fat in normal and obese mice. British Journal of Nutrition, 43(1), 83–86.

Ruegsegger, G. N., & Booth, F. W. (2018). Health benefits of exercise. Cold Spring Harbor perspectives in medicine, 8(7), a029694.

Savira, M., Rusdiana, S. S. W., & Syahputra, M. (2020). Comparison Malondialdehyde (MDA) Level between Obesity Non Metabolic Syndrome and Obesity with Metabolic Syndrome Patients. Age, 53(11.3), 44.55-10.48.

Schädel, P., Wichmann-Costaganna, M., Czapka, A., Gebert, N., Ori, A., & Werz, O. (2023). Short-Term Caloric Restriction and Subsequent Re-Feeding Compromise Liver Health and Associated Lipid Mediator Signaling in Aged Mice. Nutrients, 15(16), 3660.

Shi, Y., Zhong, L., Liu, Y., Xu, S., Dai, J., Zhang, Y., & Hu, Y. (2024). Dietary sanguinarine supplementation recovers the decrease in muscle quality and nutrient composition induced by high-fat diets of grass carp (Ctenopharyngodon idella). Animal Nutrition, 17, 208–219. 10.1016/j.aninu.2024.04.001

Shortreed, K. E., Krause, M. P., Huang, J. H., Dhanani, D., Moradi, J., Ceddia, R. B., & Hawke, T. J. (2009). Muscle-specific adaptations, impaired oxidative capacity and maintenance of contractile function characterize diet-induced obese mouse skeletal muscle. PloS one, 4(10), e7293.

Sriwijitkamol, A., Ivy, J. L., Christ-Roberts, C., DeFronzo, R. A., Mandarino, L. J., & Musi, N. (2006). LKB1-AMPK signaling in muscle from obese insulin-resistant Zucker rats and effects of training. American Journal of Physiology-Endocrinology and Metabolism, 290(5), E925–E932.

Stockwell, B. R. (2022). Ferroptosis turns 10: Emerging mechanisms, physiological functions, and therapeutic applications. cell, 185(14), 2401–2421.

Stockwell, B. R., Angeli, J. P. F., Bayir, H., Bush, A. I., Conrad, M., Dixon, S. J., Fulda, S., Gascón, S., Hatzios, S. K., & Kagan, V. E. (2017). Ferroptosis: a regulated cell death nexus linking metabolism, redox biology, and disease. cell, 171(2), 273–285.

Stockwell, B. R., & Jiang, X. (2019). A physiological function for ferroptosis in tumor suppression by the immune system. Cell Metabolism, 30(1), 14–15.

Stringer, C., Wang, T., Michaelos, M., & Pachitariu, M. (2021). Cellpose: a generalist algorithm for cellular segmentation. Nature Methods, 18(1), 100–106. 10.1038/s41592-020-01018-x

Suga, T., Kinugawa, S., Takada, S., Kadoguchi, T., Fukushima, A., Homma, T., Masaki, Y., Furihata, T., Takahashi, M., & Sobirin, M. A. (2014). Combination of exercise training and diet restriction normalizes limited exercise capacity and impaired skeletal muscle function in diet-induced diabetic mice. Endocrinology, 155(1), 68–80.

Tang, D., Chen, X., Kang, R., & Kroemer, G. (2021). Ferroptosis: molecular mechanisms and health implications. Cell Research, 31(2), 107–125. 10.1038/s41422-020-00441-1

Vellers, H. L., Letsinger, A. C., Walker, N. R., Granados, J. Z., & Lightfoot, J. T. (2017). High Fat High Sugar Diet Reduces Voluntary Wheel Running in Mice Independent of Sex Hormone Involvement. Front Physiol, 8, 628. 10.3389/fphys.2017.00628

Walsh, M. E., Shi, Y., & Van Remmen, H. (2014). The effects of dietary restriction on oxidative stress in rodents. Free Radical Biology and Medicine, 66, 88–99.

Wang, H., An, P., Xie, E., Wu, Q., Fang, X., Gao, H., Zhang, Z., Li, Y., Wang, X., & Zhang, J. (2017). Characterization of ferroptosis in murine models of hemochromatosis. Hepatology, 66(2), 449–465.

Weisberg, S. P., McCann, D., Desai, M., Rosenbaum, M., Leibel, R. L., & Ferrante, A. W. (2003). Obesity is associated with macrophage accumulation in adipose tissue. The Journal of clinical investigation, 112(12), 1796–1808.

Wenzel, S. E., Tyurina, Y. Y., Zhao, J., Croix, C. M. S., Dar, H. H., Mao, G., Tyurin, V. A., Anthonymuthu, T. S., Kapralov, A. A., & Amoscato, A. A. (2017). PEBP1 wardens ferroptosis by enabling lipoxygenase generation of lipid death signals. cell, 171(3), 628-641. e626.

Xiang, Y.-Y., Baek, K.-W., Won, J.-H., Park, Y., Kim, J.-S., Xiang, Y.-Y., Baek, K.-W., Won, J.-H., Park, Y., & Kim, J.-S. (2023). Effects of Lifelong Aerobic Exercise on Ferroptosis-Related Gene Expressions in Kidney of Aged Mice. Exercise Science, 32(4), 410–418.

Yang, W. S., SriRamaratnam, R., Welsch, M. E., Shimada, K., Skouta, R., Viswanathan, V. S., Cheah, J. H., Clemons, P. A., Shamji, A. F., & Clish, C. B. (2014). Regulation of ferroptotic cancer cell death by GPX4. cell, 156(1), 317–331.

Yang, Y., Smith, D. L., Jr., Keating, K. D., Allison, D. B., & Nagy, T. R. (2014). Variations in body weight, food intake and body composition after long-term high-fat diet feeding in C57BL/6J mice. Obesity (Silver Spring), 22(10), 2147–2155. 10.1002/oby.20811

Yang, Y., Smith Jr, D. L., Keating, K. D., Allison, D. B., & Nagy, T. R. (2014). Variations in body weight, food intake and body composition after long-term high-fat diet feeding in C57BL/6J mice. Obesity, 22(10), 2147–2155.

Yeu, J., Ko, H. J., Kim, D., Ahn, Y., Kim, J., Lee, W., Jung, I., Suh, J., & Lee, S. J. (2019). Evaluation of iNSiGHT VET DXA (Dual-Energy X-ray Absorptiometry) for assessing body composition in obese rats fed with high fat diet: a follow-up study of diet induced obesity model for 8 weeks. Lab Anim Res, 35, 2. 10.1186/s42826-019-0004-2

Yu, Y., Jiang, L., Wang, H., Shen, Z., Cheng, Q., Zhang, P., Wang, J., Wu, Q., Fang, X., & Duan, L. (2020). Hepatic transferrin plays a role in systemic iron homeostasis and liver ferroptosis. Blood, The Journal of the American Society of Hematology, 136(6), 726–739.

Zhang, X., Ma, Y., Lv, G., & Wang, H. (2023). Ferroptosis as a therapeutic target for inflammation-related intestinal diseases. Front Pharmacol, 14, 1095366. 10.3389/fphar.2023.1095366

Zhang, Y., Fischer, K. E., Soto, V., Liu, Y., Sosnowska, D., Richardson, A., & Salmon, A. B. (2015). Obesity-induced oxidative stress, accelerated functional decline with age and increased mortality in mice. Archives of biochemistry and biophysics, 576, 39–48.

Zhao, X., Si, L., Bian, J., Pan, C., Guo, W., Qin, P., Zhu, W., Xia, Y., Zhang, Q., & Wei, K. (2022). Adipose tissue macrophage-derived exosomes induce ferroptosis via glutathione synthesis inhibition by targeting SLC7A11 in obesity-induced cardiac injury. Free Radical Biology and Medicine, 182, 232–245.

Zhuang, A., Yang, C., Liu, Y., Tan, Y., Bond, S. T., Walker, S., Sikora, T., Laskowski, A., Sharma, A., de Haan, J. B., Meikle, P. J., Shimizu, T., Coughlan, M. T., Calkin, A. C., & Drew, B. G. (2021). SOD2 in skeletal muscle: New insights from an inducible deletion model. Redox Biol, 47, 102135. 10.1016/j.redox.2021.102135

